# Effect of Alpha-Lactalbumin and Lactoferrin Oleic Acid Complexes on Chromatin Structural Organization

**DOI:** 10.1101/675223

**Authors:** Dmitry V Lebedev, Yana A Zabrodskaya, Vitaly Pipich, Alexander I Kuklin, Edward Ramsay, Alexey V Sokolov, Anna Yu Elizarova, Aram A Shaldzhyan, Natalia A Grudinina, Rimma A Pantina, Baohu Wu, Tatiana A Shtam, Andrey V Volnitskiy, Alexander E Shmidt, Alexey V Shvetsov, Vladimir V Isaev-Ivanov, Vladimir V Egorov

## Abstract

This work focuses on the study of multimeric alpha-lactalbumin oleic acid and lactoferrin oleic acid complexes. The purpose of the research is to study possible mechanisms involved in their pro-apoptotic activities, as seen in some tumor cell cultures. Complexes featuring oleic acid (OA) with human alpha-lactalbumin (hAl) or with bovine alpha-lactalbumin (bAl), and human lactoferrin (hLf) were investigated using small-angle neutron scattering (SANS). It was shown that while alpha-lactalbumin protein complexes were formed on the surface of polydisperse OA micelles, the lactoferrin complexes comprised a monodisperse system of nanoscale particles. Both hAl and hLf complexes appeared to interact with the chromatin of isolated nuclei affecting chromatin structural organization. The possible roles of these processes in the specific anti-tumor activity of these complexes are discussed.

## Introduction

Cancer is the second most common cause of death in the world after cardiovascular diseases [1]. Targeting tumor cells is one of the key problems in developing anti-cancer therapies. Generally speaking, there are two approaches to the targeting of cancerous cells. In the first variant, agents are developed which are broadly toxic, and subsequent development work is completed which aims to ensure that such agents are delivered only to cancer cells. In the second variant, the delivery of agents is rather broad, but the agent itself is known to be almost exclusively toxic to cancer cells. Here, we focus on complexes which fall into the second category. Human alpha-lactalbumin (hAl, 14.4 kDa) is a whey protein and, normally, it is only in the mammary gland during lactation. The main function of alpha-lactalbumin is alteration of galactosyltransferase substrate specificity. Alpha-lactalbumin is expressed in many breast tumors [2].

Published work indicates that multimeric complexes of lactalbumin and oleic acid (CAlOA) have antitumor activity *in cellulo*, yet without toxicity to normal cells. [3]. The mechanism of action behind such complexes is not well understood. It was shown that CAlOA is capable of penetrating into cells [4], interacting with chromatin [5,6], and interacting with nucleotide-binding proteins [7].

In our previous work, in which we prepared CAlOA complexes [8], it was shown that they had similar activities to those described in the literature for HAMLET (Human Alpha-lactalbumin Make LEthal to Tumor cells) complexes [9]. It should be noted that CAlOA differs from HAMLET in terms of preparation and, most likely, in terms of structure. In recent years it has become obvious that hAl is not the only milk protein capable of binding oleic acid. Lactoferrin (Lf, 78 kDa), for example, also forms cytotoxic complexes with OA [10].

The aim of this research is to study the structure of OA/human alpha-lactalbumin (ChAlOA) and OA/bovine alpha-lactalbumin (CbAlOA) complexes and to determine their influences on tumor chromatin structure, including understanding potential antitumor mechanisms. The properties of ChLfOA (human lactoferrin in complex with OA) were also studied.

## Experimental

### Materials

Commercial oleic acid, reagents, buffers, and chromatography column sorbents were purchased from Merck (Germany).

### Alpha-lactalbumin purification

Human and bovine alpha-lactalbumin (hAl and bAl, respectively) were isolated from fresh human or bovine milk by hydrophobic chromatography using phenyl-sepharose, as described [11]. Protein purities and identities were confirmed by SDS-PAGE [12], followed by Coomassie staining [13] and MALDI mass spectrometric analysis.

### Human lactoferrin purification

Lactoferrin from breast milk (hLf) was isolated using ion exchange and gel-filtration chromatography, as described [14]. Briefly, breast milk was centrifuged twice (500*g*, 4°C, 1 hour; and 10,000*g*, 4°C, 30 minutes) to remove lipids. Supernatant was filtered, centrifuged at 10,000*g* for 30 minutes, and filtered again. Skimmed milk was applied to a column (5×12 cm) containing CM-Sephadex (Sigma-Aldrich) and washed by phosphate buffered saline (PBS). Human lactoferrin was next eluted by linear NaCl gradient (0.15 M – 1 M) containing 10 mM sodium phosphate buffer (pH 7.4). At the next stage, hLf was further purified by gel-filtration chromatography using Sephacryl S-200 HR. The protein purity and identity were confirmed by SDS-PAGE [12] followed by Coomassie staining [13] and MALDI mass spectrometric analysis.

### Mass spectrometry

The amino acid sequences of the isolated proteins (hAl, bAl, hLf) were verified by MALDI mass spectrometry. Protein bands after SDS-PAGE were excised and washed, as described [15] without reduction or modification of disulphide bonds. Samples subjected to tryptic digestion with sequence grade modified porcine trypsin (Promega, USA) for 5 hours, 37°С. The spectra of resultant tryptic peptides were measured using an HCCA matrix (Bruker, Germany) and the UltrafleXtreme MALDI-TOF/TOF mass spectrometer (Bruker, Germany) in positive ion registration mode. Identification was performed using MASCOT (matrixscience.com) and the NCBI database (ncbi.nlm.nih.gov). Accuracy was limited to 20 ppm. Methionine oxidation was selected as variable modification. Protein identification was considered reliable when Scores exceeded the threshold value (p < 0.05).

### OA/human alpha-lactalbumin (ChAlOA) and OA/bovine alpha-lactalbumin (CbAlOA) complexes

To prepare complexes, oleic acid, in excess, (500 ul) was added to 500 ul of alpha-lactalbumin solution (5 mg/ml in PBS), and mixed vigorously using an ultrasonic bath (Branson, USA) for 5 minutes. After centrifugation (12,000*g*, 4ºC, 10 min.), the lower fraction containing pure complexes was carefully collected.

### Oleic acid/human lactoferrin complexes (ChLfOA)

A three layer approach was used to prepare ChLfOA. The first (bottom) layer was a 4 ml hLf solution (160 mg/ml in PBS). One milliliter of PBS was carefully layered onto the base solution layer, followed by addition of 100 ul of ethanol; the ethanol was carefully added to the top (PBS) fraction in order to avoid direct contact with protein solution. To prepare oleic acid solution 17 ul of OA was mixed with 500 ul of ethanol which correspond 8-fold molar excess of OA to hLf. Complexes were obtained by hLf titration with aliquots of OA solution (25 ul) which is followed by throughly mixing tree times with 1 minute interval. Solution of hLf and OA mixture remains clear while PBS with OA is cloudy. ChLfOA in ethanol was dialyzed against PBS and filtered (0.22 um). Ration of hLf and OA in complexes as 1 to 8 was confirmed using colorimetric kit for nonesterified fatty acids detection (Randox, F115).

### HeLa nuclei isolation

Was performed as described [16]. Isolated nuclei were treated with ChAlOA or ChLfOA (0.5 mg/ml) for 15 minutes, washed with PBS and fixed by 0.5% gluthar aldehyde.

### Small angle neutron scattering (SANS)

For SANS experiments all samples were transferred in PBS dissolved on D_2_O (PBS/D_2_O).

CbAlOA and bAl spectra were obtained with the YuMO spectrometer located on the fourth channel of the IBR-2 high-flux reactor (Frank Laboratory of Neutron Physics, International Intergovernmental Organization Joint Institute for Nuclear Research, Dubna, Russia). Measurements were performed in standard geometry, described in [17,18], using two detectors simultaneously in the range of *q = 4π/λ sin θ/2* from 0.006 to 0.3 Å^−1^. Data were preliminary processed as described [19]: the normalization and obtaining of curves in absolute units is realized using the metallic vanadium located directly in front of the detector.

ChAlOA and ChLfOA spectra were obtained with the KWS2 spectrometer (Jülich Centre for Neutron Science at Heinz Maier-Leibnitz Zentrum, FRM-II reactor, Munich, Germany) in the *q* region from 7.5×10^−3^ to 0.25 Å^−1^ using two position of detector.

Spectra of isolated HeLa nuclei treated with ChAlOA or ChLfOA were obtained with the KWS2 spectrometer (two position of detector) and KWS3 (MLZ, JCNS, Munich, Germany) in the *q* region from 10^−4^ to 0.25 Å^−1^.

All curves were analyzed using Origin2015 software.

## Results and Discussion

All isolated proteins which were used for complexes preparation were reliable identified by mass spectrometry as bAl, hAl and hLf respectively. To characterize complexes of alpha-lactalbumin with oleic acid, CbAlOA and ChAlOA were studied by SANS; similar solutions of bAl and pure OA were used as control (Figure 1). The data are presented on the double-logarithmic scale.

**Figure 1.**
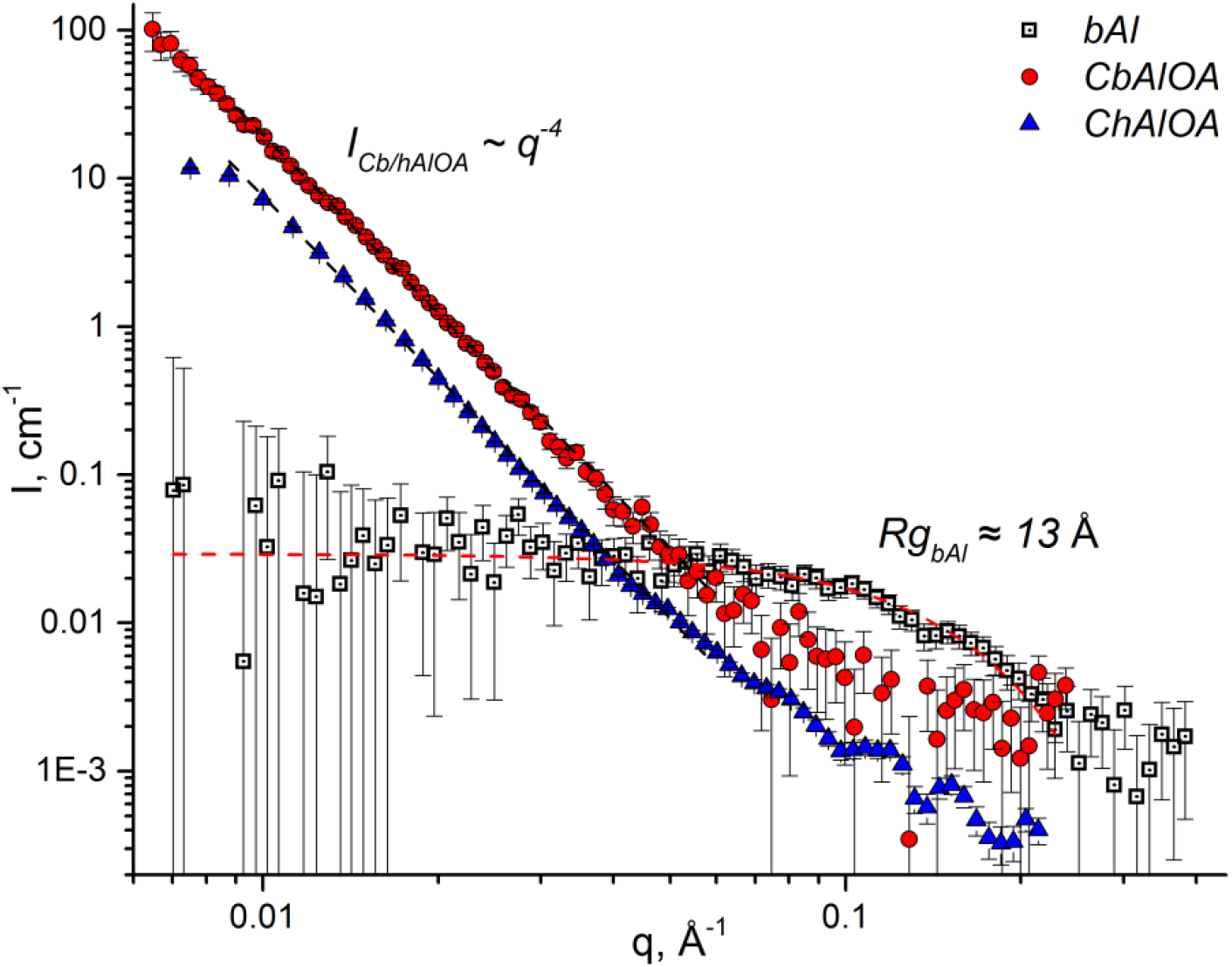
SANS spectra of bAl (squares), CbAlOA (circles) and ChAlOA (triangles). Scattering of OA coincided with buffer (not shown).

*I* – scattering intensity,
*q* – magnitude of the momentum transfer,
*Rg* – gyration radius.

Samples of CAlOA were characterized by high differential cross section of scattering in small angles that decayed away by Porod’s law (*1/q^4^*). Data indicates that scattering in the sample occurs on the surface of big particles (dozens of nanometers and more), that are presumably represented by stable suspension of OA in PBS/D_2_O. At the same time SANS spectrum of pure bAl follows the Guinier law (*Rg^2^/3* ~ *ln(I(q))/q^2^*) with *Rg* ~ 13 Å. It should be noted that scattering intensity of analogous sample of OA without bAl coincided with buffer (data not shown). It could be concluded that bAl is necessary for OA suspension formation. Micelles formed in presence of bAl are stable for a few days.

CbAlOA and ChAlOA were prepared analogously to those that we studied earlier [8]. It was shown that such complexes have the characteristic for ChAlOA antimicrobial activity. In the same study complexes were characterized by atomic force microscopy and chromatography. In the present research CAlOA structure was studied by SANS method, which is sensitive to OA particles and have high contrast in D2O containing buffer.

In reference [20] using small angle X-ray scattering (SAXS) it was shown that small lactalbumin and oleic acid complexes are present in the ChAlOA solution; in the later work of the same research group, performed using SANS, the presence of larger (few nanometers) drops of OA was shown in the center of the scattering objects [21]. In our experiment we observe CAlOA to contain large (tens or hundreds of nanometers) drops of OA.

In contrast to alpha-lactalbumin complexes, the SANS spectra of ChLfOA, shown in Figure 2, indicate the absence of noticeable amounts of OA micelles and the presence of ChLfOA complexes that are significantly larger (*Rg* ~ 8 nm) than lactoferrin itself (~ 5 nm).

**Figure 2.**
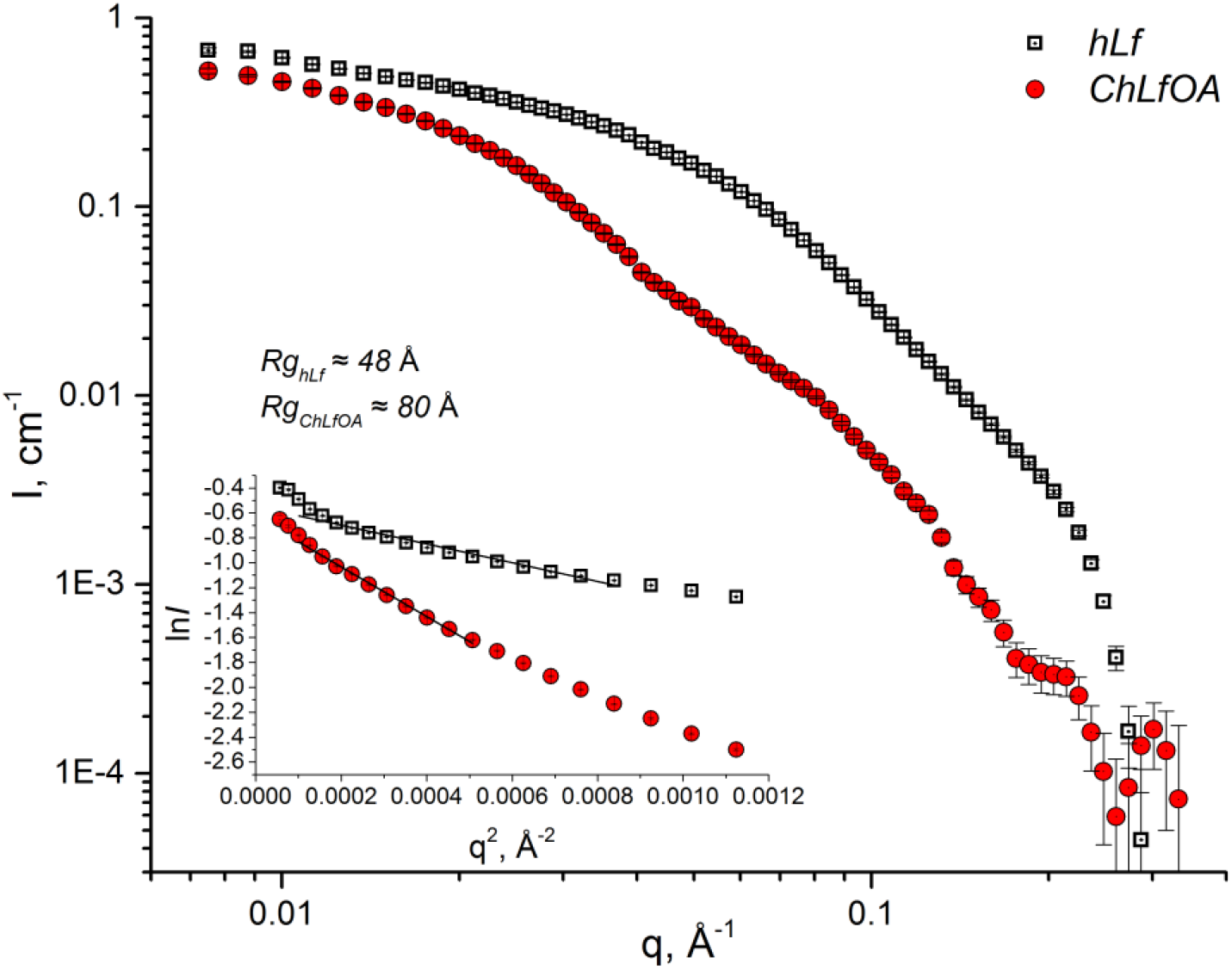
The SANS spectra of hLf (squares) and ChLfOA (circles).

*I* – scattering intensity,
*q* – magnitude of the momentum transfer,
*Rg* – gyration radius. *Rg* is calculated from the slope of linear fit of plot presented in Guinier scale (ln*I* vs. q^2^) using the equation *Rg*^2^/3 ~ *ln(I(q))/q*^2^.

Thus, large micelles of oleic acid were detected in the samples of CAlOA, but not ChLfOA. At the same time we cannot exclude the presence of smaller particles of the ChAlOA complex, that cannot be characterized due to the large background of scattering on oleic acid. The particular components (OA micelles or true protein-OA complexes) of the sample that underly the previously observed [8] biological activity of CAlOA are still to be established. The exclusive role of OA in protein-OA complexes activity and absence of the role of the proteins themselves were hypothesized in [22]. On the other hand, there were no detectable amount of micelles in ChLFOA, and its biological activity [23] should be most likely be attributed to true multimolecular complexes.

We hypothesized that complexes of Al or Lf with OA influence on chromatin structure when penetrating cell. Accordingly, we have studied the effect of ChAlOA and ChLfOA on HeLa cells in model system – isolated HeLa nuclei. The HeLa nuclei curves (Figure 3, squares) has two phases fractal nature (with power law exponent D ~ 3.5 and ~ 2.5) with crossover point (conforming to the transition from the three-dimensional to the surface fractal) similar to those observed previously for the chromatin nuclei of a variety of cells [24] [25].

**Figure 3.**
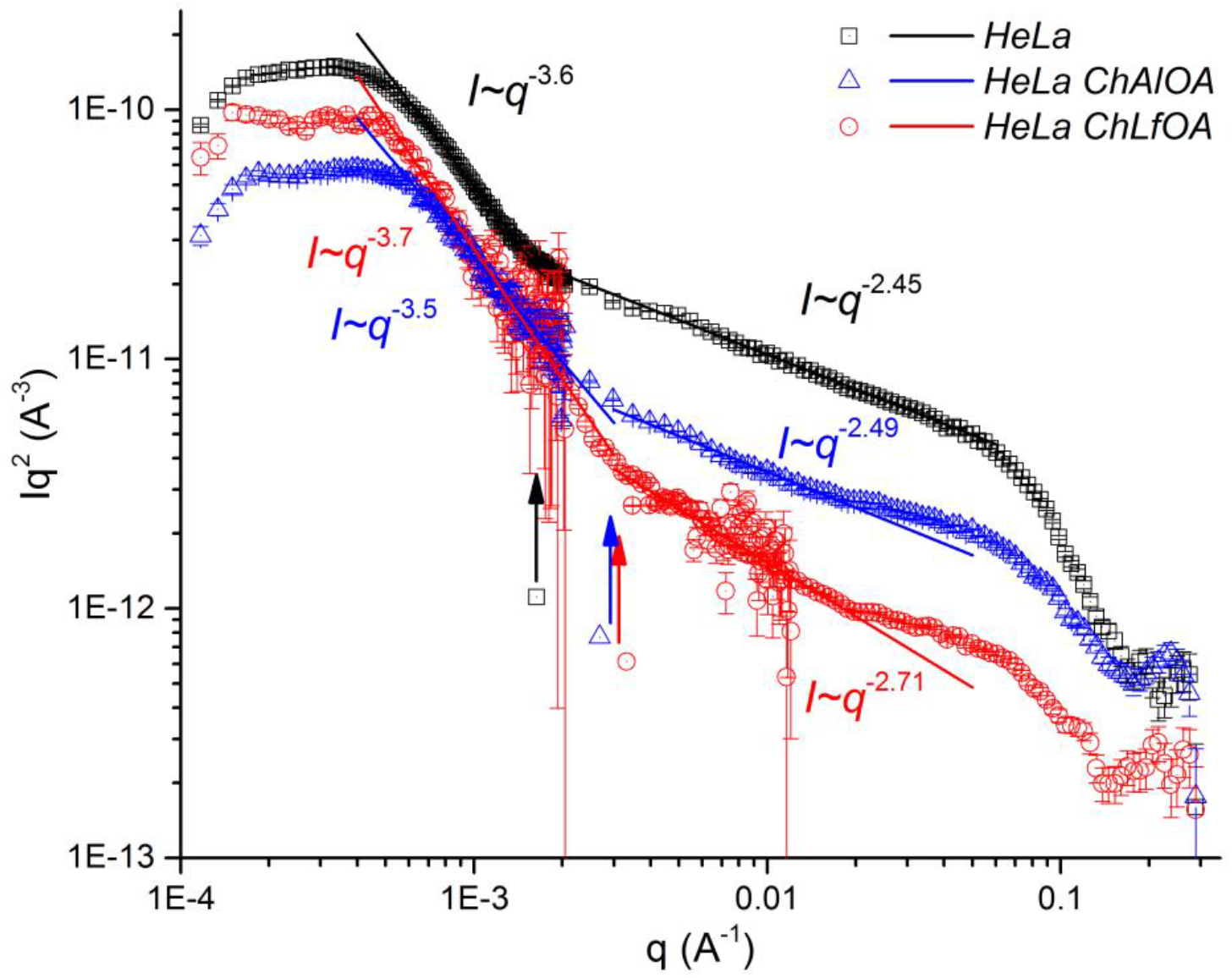
SANS spectra (double-logarithmic Kratky scale representation) of isolated HeLa nuclei: mock treated (squares) and treated with ChAlOA (triangles) or ChLfOA (circles). Data from both KWS-2 and KWS-3 spectrometers shown on the same plot. Arrows mark the approximate crossover points on HeLa, HeLa ChAlOA and HeLa ChLfOA curves.

Isolated HeLa nuclei were treated with ChAlOA or ChLfOA followed by SANS spectra registration (Figure 3, triangles and circles respectively). It was observed that curves demonstrate some changes in *q* region corresponding to short-range nucleosome structure (0.04 – 0.08 Å^−1^), as well as in fractal characteristics of large-scale chromatin structure. Interestingly, the crossover point tends to move under the influence of ChAlOA and ChLfOA from ~ 400 nm (for mock treated HeLa nuclei) to ~ 200 nm (for treated HeLa nuclei). Based on the nature of changes (increase in fractal dimension in *q* range 0.002 – 0.02 Å^−1^) we proposed that complexes studied cause chromatin compaction. In a cell, this can lead to a decrease in the expression of genes in compacted regions, and thus interfere with the functioning of tumor cells. It should be taken into account that the addition of hAl or hLf in the absence of OA as well as addition of protein-OA complexes treated with detergent (Triton X-100) did not lead to changes in the chromatin structure detected by SANS (data not shown).

Thus, the effect of ChLfOA on the chromatin structure in HeLa nuclei not only qualitatively coincide with the ChAlOA effect but is more pronounced. This suggests that the presence of OA micelles or OA molecules is not the main mechanism of the effect of ChLaOA or ChLfOA complexes on chromatin organization.

## Conclusion

Thus, the artificially created complex of multimeric alpha-lactalbumin, formed in the presence of oleic acid, contain a stable suspension of oleic acid with particles of various sizes, up to tens, and possibly hundreds of nanometers. Alpha-lactalbumin molecules, presumably, are localized on the surface of such particles. At the same time, the lactoferrin and oleic acid complex, does not contain large micelles, and can be considered as a monodisperse system of small (~ 20 nm) particles. When exposed to isolated HeLa nuclei, both ChAlOA and ChLfOA lead to a change in the chromatin structure, which can determine their antitumor effect.

## Acknowledgements

This work based on experiments provided on KWS-3 (#11603, #13835) and KWS-2 (#11589) instruments, local contacts: V. Pipich and A. Radulesku, MLZ, FRM- II Munich, Germany, and also on YuMO spectrometer (experiments #2015-10-15-17-15-09, #2015-10-15-19-57-02, #2017-04-14-14-35-32), IBR-2, local contact A.I. Kuklin, Dubna, Russia.

